# The link between supplementary tannin level and conjugated linoleic acid (CLA) formation in ruminants: A meta-analysis

**DOI:** 10.1101/612523

**Authors:** Rayudika Aprilia Patindra Purba, Pramote Paengkoum, Siwaporn Paengkoum

## Abstract

This meta-analysis was conducted to predict and assert a way to discover conjugated linoleic acid (CLA) formation in ruminant-derived products as problem solver of human health issues threated by plant-containing tannins. The objective was to expound, to compare, and to confirm the efficiency of tannins cultivating CLA formation whether using *in vitro* and/or *in vivo* study. A database was created using the ruminants with selectively 26 experiments comprising 683 dietary treatments as explained *in vitro* and *in vivo* methods that were applied as a statistical SAS 9.4 tool. Basically, increasing level of tannins leaded to an underlying decrease in CLA formation (p<0.001), initially at predicting coefficient determination R^2^=0.193, R^2^=0.929, and R^2^=0.549 for CLA *in vitro, in vivo* of CLA milk shift, and *in vivo* of CLA meat precipitation, respectively. *In vitro* may accurately predict to the *in vivo* observation. Unfortunately, there were no relationship *in vitro* towards *in vivo* observation (R^2^<0.1). It indicated to be difficult to predict CLA from *in vitro* to *in vivo* separately situations. According to all studies, the level of tannin’s utilization for inhibiting biohydrogenation was not exceedingly >50 g/kg DM recommended. Secondly, the *in vivo* method was more suitable for directly observation that concerned in fatty acid transformation.

## Introduction

Nowadays, the consumers have been aware to more selectively in their consumption, especially ruminant-derived products such a concerning fat composition in milk and meat. Lourenço, et al [1] reviewed that food for human derived from ruminant product is a high of Saturated Fatty Acid (SFA) and has lower polyunsaturated fatty acid (PUFA) due to detrimental condition of human health, including intensified serum low-density lipoprotein (LDL) cholesterol level, which is a risk factor for coronary heart disease. In previous studies, have been coined conjugated linoleic acid (CLA) as natural fatty acid (FA) and this FA could solve aforementioned human problems [1-3]. The predominant isomer of CLA is cis-9, trans-11 18:2, representing 75–90% of the total CLA in ruminant fat, and trans-7, cis-9 CLA is the second most prevalent isomer at 3–16% of the total CLA [1, 4] and the trans-11, 18:1 (vaccenic acid) existence is notable know to support cis-9, trans-11 18:2 [3]. However, producing CLA in milk and meat is quite difficult because its process invites biohydrogenation respecting to catalyzation by ruminal microorganisms. For instance, *Butyrifibrio fibrisolvens* was identified to undertake biohydrogenation of FA and to carry in creating cis-9, trans-11 18:2 and trans-11 18:1 by way of trans-11 18:2 (n-6) [5, 6]. Thus, bacteria acts the fundamental role in FA biohydrogenation [7] and looking for alternative feed additives from Phytochemicals [8] as anti-microbial could be greater option to increase CLA in ruminant products.

Essential oils are commonly supplementation derived from plant and marine product. Their function exactly had many modes bringing ruminal bacteria N down [9] and inhibited survival of *Butivibrio fibrisolvens* and *Butivibro proteoclasicus* community on biohydrogenation[10]. The effective of essential oils had variable impacts on ruminal fermentation [11], it might be believed in depend on source extraction, method, dose, basic diet, pH and preliminary period of microorganism to adapt essential oils. Subsequently, forage feeding is used gaining long chain of PUFA as galacto-, sulfo-, and phospholipid that could exert keeping PUFA long time on biohydrogenation. Besides, ionosphere feed additive namely saponins and tannins is coming to deserve attention as antimicrobial properties. Li, et al [12], ability of tannin supplementation had a broadly distortion of rumen microbiota, thereby being useful to shift rumen performance. Another, saponins concerned to inhibit methane emission and lower biohydrogenation because of defaunation function leading to protozoa-lipid population decrease [9]. On other hand, tannins had a greater impressive mode, particularly antimicrobial behavior to assert *Clostridium proteoclasticus* converting trans-11, 18:1 to 18:0 form [10]. Consequently, tannins could be appreciated as a temporary fraction to improve CLA production in FA composed manipulation of ruminal fermentation.

Tannins including condensed and hydrolyzed forms have expressed widely antimicrobial properties in rumen studies. In previous years, Jayanegara, et al [13], conducted meta-analysis with collecting data from *in vitro* and *in vivo* studies that supplementing feed-containing tannins in rumen feedstuff diminished methane level and affected to palatability of ruminant. Also, Jerónimo, et al [14], reviewed chemical structure of tannins behaved rough effect on animal performance and the quality of their products (meat and milk) particularly on the fatty acid profile, oxidative stability, and organoleptic properties. In these two publications explained valuable tannins, edible usages, and potential functions separately. Although, no one even in single chapter addressed a relationship of tannin supplementation in rumen diet towards to biohydrogenation approaching with the meta-analysis technique. The clear-cut method whether using *in vitro* and/or *in vivo* to provide a prerequisite is also needed. Hopefully, the result of present study could be useful for animal science, animal nutrition, and biotechnology expertise. Therefore, the objectives were (i) to expound the effectiveness of tannins modulating CLA formation, (ii) to study comparison of gained result based on *in vitro* and *in vivo* methods, (iii) to confirm the relationship between *in vitro* and *in vivo* studies applying the meta-analysis as a statistical tool.

## Methods

### Search strategy and selection criteria

A database created from experiments which the dietary tannin concentrations and CLA properties were concerned touching closer to PRISMA (Preferred Reporting Items for Systematic Reviews and Meta-Analyses) [15], see Fig 1. These data were gathered on the ISI Web of Science (recently form in ISI Web of Knowledge) database using “conjugated linoleic acid,” “biohydrogenation”, “rumen,” “tannin,” “meat,” “milk,” “*in vivo*,” and “*in vitro*” as keywords from October 31, 2016 to March 23, 2019. Title/abstract, topic and keyword search terms were used in combination. Results were limited to trials published in English (S1 Table). For further consideration, the results were touched with single search in relevant studies and reviews. Endnote (Thompson ISI Research-Soft, Philadelphia, PA, US) was used to repository the relevant articles and remove duplicate articles.

**Fig 1.**
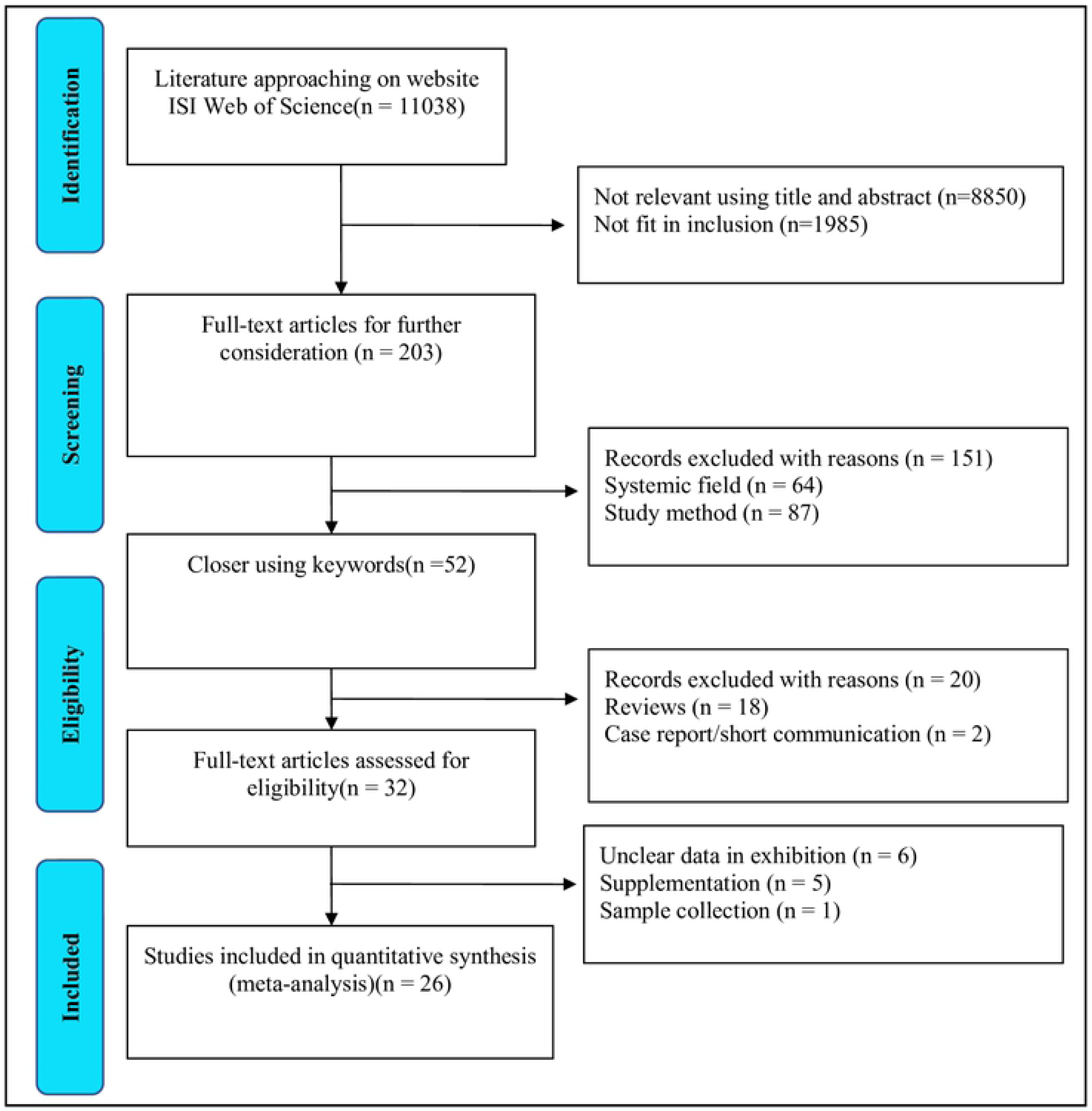
Modified flow chart of the selection process for the eligible studies.

### Study criteria, quality assessment, and data extraction

Studies were included if they met the following criteria: (1) the study design was an *in vitro*; (2) the study design was an *in vivo*; (3) Object used ruminants: cow, goat, sheep in dairy or meat product (4) relevant data was retrievable; and (5) the studies were published after 1 December 2008. Authors were contacted by e-mail and ResearchGate provider, if data had questionable. If that was unsuccessful, references were excluded on account of inaccessibility of data.

The raw data were strictly screened and accepted in similar calculating unit per parameter, e.g., g/kg FAME (fatty acid methyl ester) and g/kg DM (dry matter) for all FA and tannin level, respectively. Finally, the comprehensive database consisted of 683 dietary treatments in 26 experiments as explanation in Table 1 (*in vitro* experiments) and Table 2 (*in vivo* experiments). In addition, the sources were collected even deriving from individually publication. The database was picked selectively into two categories based on different methods or systems applied in the experiments. As a result, there were *in vitro* batch culture (10 experiments/356 treatments) and *in vivo* experiments (16/327) with 10 experiments concerning CLA level from milk source and 6 others from intra muscular fatty acid in meat source, completing with 2 experiments conducted both *in vitro* and *in vivo* on the same time. The Cochrane Reviewer’s Handbook 4.2 was used to assess the risk of bias.

**Table 1.**
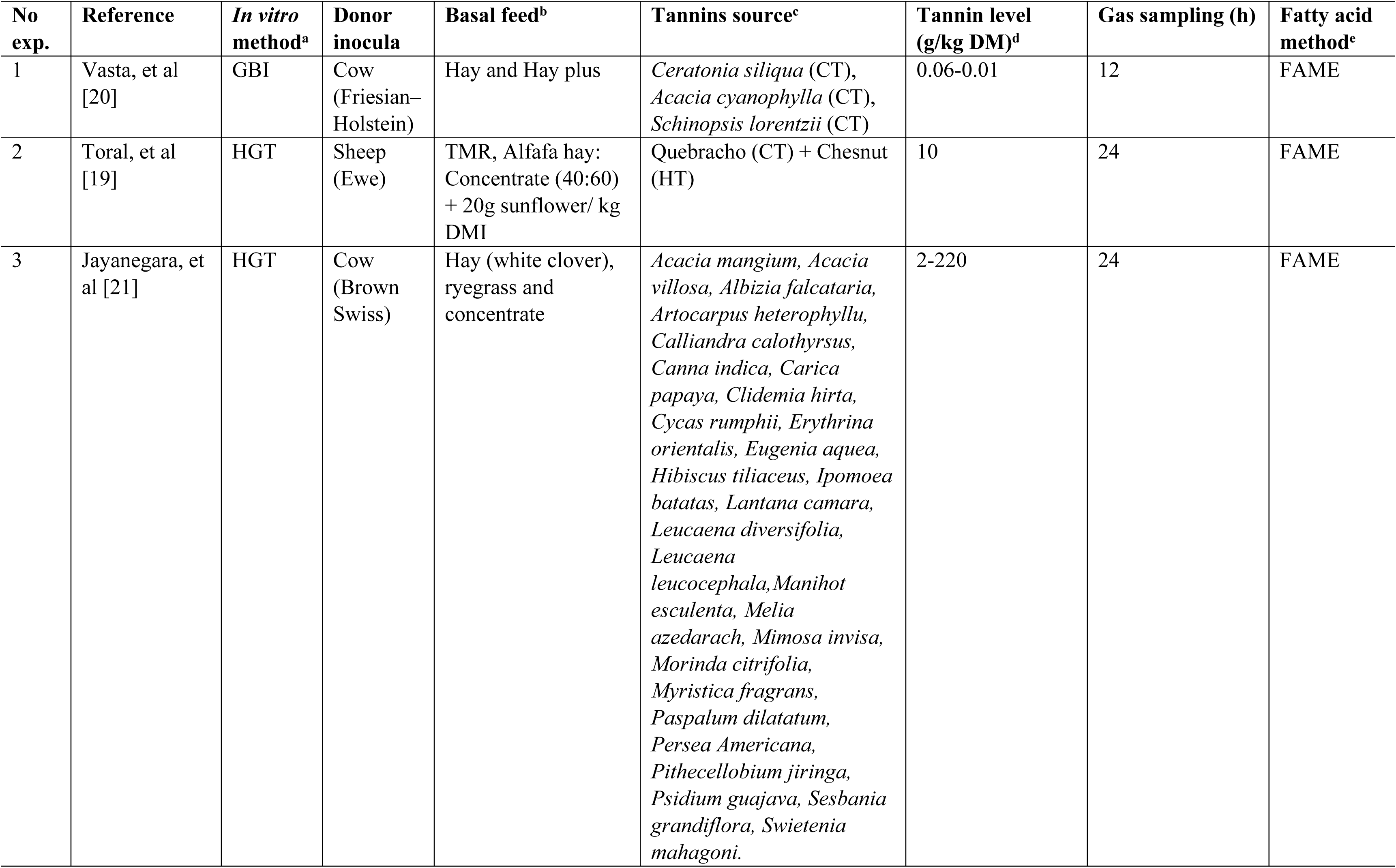
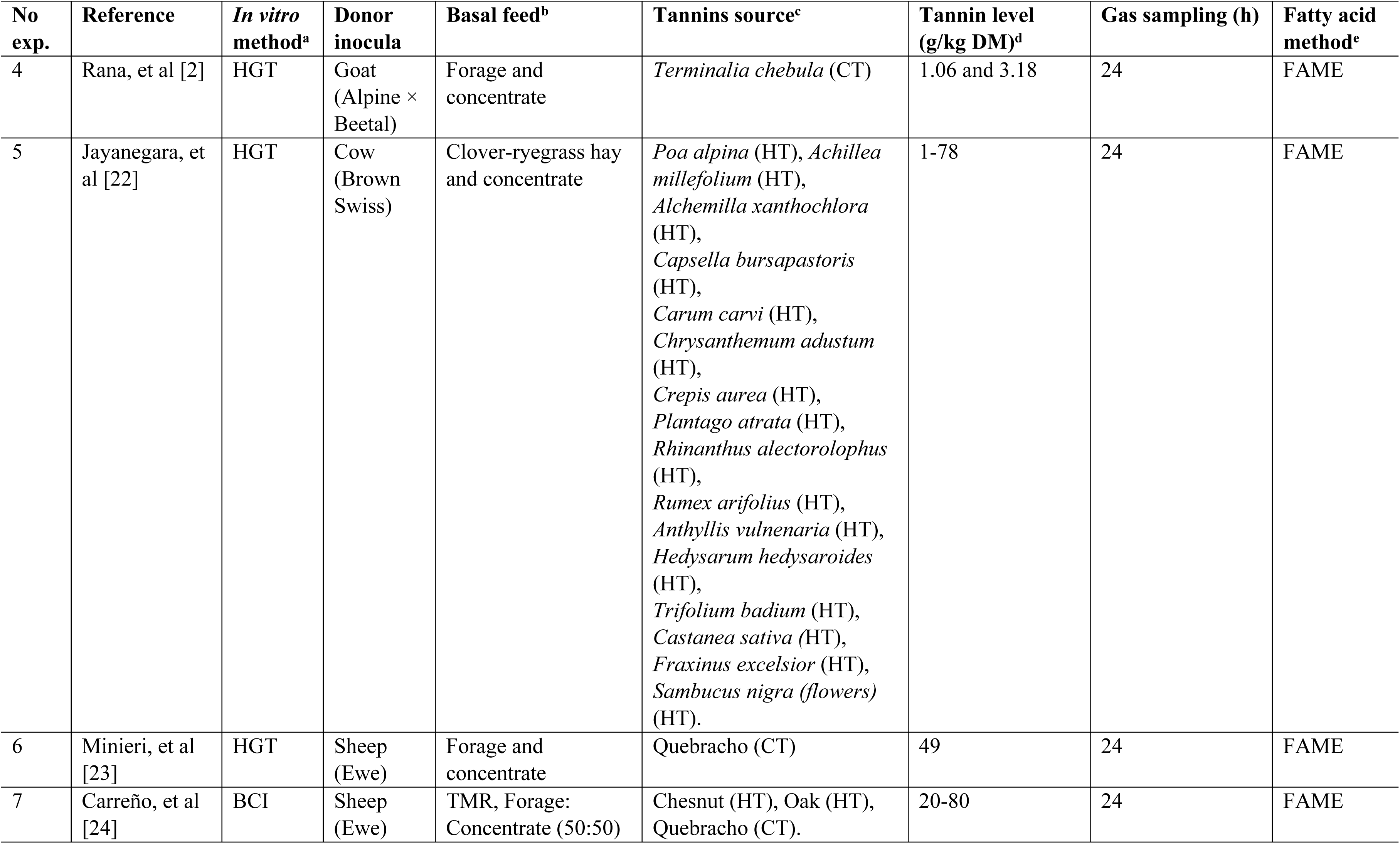
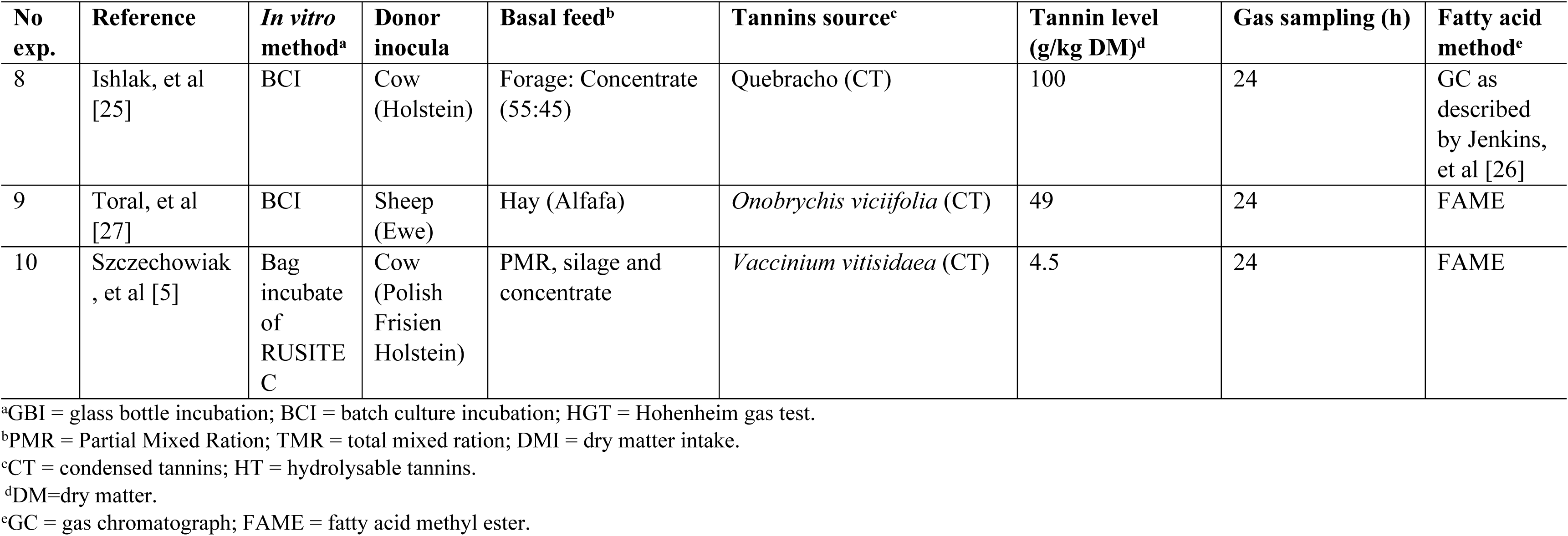
Data tabulation of i*n vitro* experiments.

| No exp. | Reference | *In vitro* method<sup>a</sup> | Donor inocula | Basal feed<sup>b</sup> | Tannins source<sup>c</sup> | Tannin level (g/kg DM)<sup>d</sup> | Gas sampling (h) | Fatty acid method<sup>e</sup> |
| --- | --- | --- | --- | --- | --- | --- | --- | --- |
| 1 | Vasta, et al [20] | GBI | Cow (Friesian–Holstein) | Hay and Hay plus | *Ceratonia siliqua* (CT),<br>*Acacia cyanophylla* (CT),<br>*Schinopsis lorentzii* (CT) | 0.06-0.01 | 12 | FAME |
| 2 | Toral, et al [19] | HGT | Sheep (Ewe) | TMR, Alfafa hay: Concentrate (40:60) + 20g sunflower/ kg DMI | Quebracho (CT) + Chesnut (HT) | 10 | 24 | FAME |
| 3 | Jayanegara, et al [21] | HGT | Cow (Brown Swiss) | Hay (white clover), ryegrass and concentrate | *Acacia mangium*, *Acacia villosa*, *Albizia falcataria*, *Artocarpus heterophyllu*, *Calliandra calothyrsus*, *Canna indica*, *Carica papaya*, *Clidemia hirta*, *Cycas rumphii*, *Erythrina orientalis*, *Eugenia aquea*, *Hibiscus tiliaceus*, *Ipomoea batatas*, *Lantana camara*, *Leucaena diversifolia*, *Leucaena leucocephala*, *Manihot esculenta*, *Melia azedarach*, *Mimosa invisa*, *Morinda citrifolia*, *Myristica fragrans*, *Paspalum dilatatum*, *Persea Americana*, *Pithecellobium jiringa*, *Psidium guajava*, *Sesbania grandiflora*, *Swietenia mahagoni*. | 2-220 | 24 | FAME |
| 4 | Rana, et al [2] | HGT | Goat (Alpine × Beetal) | Forage and concentrate | *Terminalia chebula* (CT) | 1.06 and 3.18 | 24 | FAME |
| 5 | Jayanegara, et al [22] | HGT | Cow (Brown Swiss) | Clover-ryegrass hay and concentrate | *Poa alpina* (HT), *Achillea millefolium* (HT), *Alchemilla xanthochlora* (HT), *Capsella bursapastoris* (HT), *Carum carvi* (HT), *Chrysanthemum adustum* (HT), *Crepis aurea* (HT), *Plantago atrata* (HT), *Rhinanthus alectorolophus* (HT), *Rumex arifolius* (HT), *Anthyllis vulneraria* (HT), *Hedysarum hedysaroides* (HT), *Trifolium badium* (HT), *Castanea sativa* (HT), *Fraxinus excelsior* (HT), *Sambucus nigra* (flowers) (HT). | 1-78 | 24 | FAME |
| 6 | Minieri, et al [23] | HGT | Sheep (Ewe) | Forage and concentrate | Quebracho (CT) | 49 | 24 | FAME |
| 7 | Carreño, et al [24] | BCI | Sheep (Ewe) | TMR, Forage: Concentrate (50:50) | Chesnut (HT), Oak (HT), Quebracho (CT). | 20-80 | 24 | FAME |
| 8 | Ishlak, et al [25] | BCI | Cow (Holstein) | Forage: Concentrate (55:45) | Quebracho (CT) | 100 | 24 | GC as described by Jenkins, et al [26] |
| 9 | Toral, et al [27] | BCI | Sheep (Ewe) | Hay (Alfafa) | *Onobrychis viciifolia* (CT) | 49 | 24 | FAME |
| 10 | Szczechowiak, et al [5] | Bag incubate of RUSITE C | Cow (Polish Frisien Holstein) | PMR, silage and concentrate | *Vaccinium vitisidaea* (CT) | 4.5 | 24 | FAME |

<sup>a</sup>GBI = glass bottle incubation; BCI = batch culture incubation; HGT = Hohenheim gas test.
<sup>b</sup>PMR = Partial Mixed Ration; TMR = total mixed ration; DMI = dry matter intake.
<sup>c</sup>CT = condensed tannins; HT = hydrolysable tannins.
<sup>d</sup>DM=dry matter.
<sup>e</sup>GC = gas chromatograph; FAME = fatty acid methyl ester.

**Table 2.**
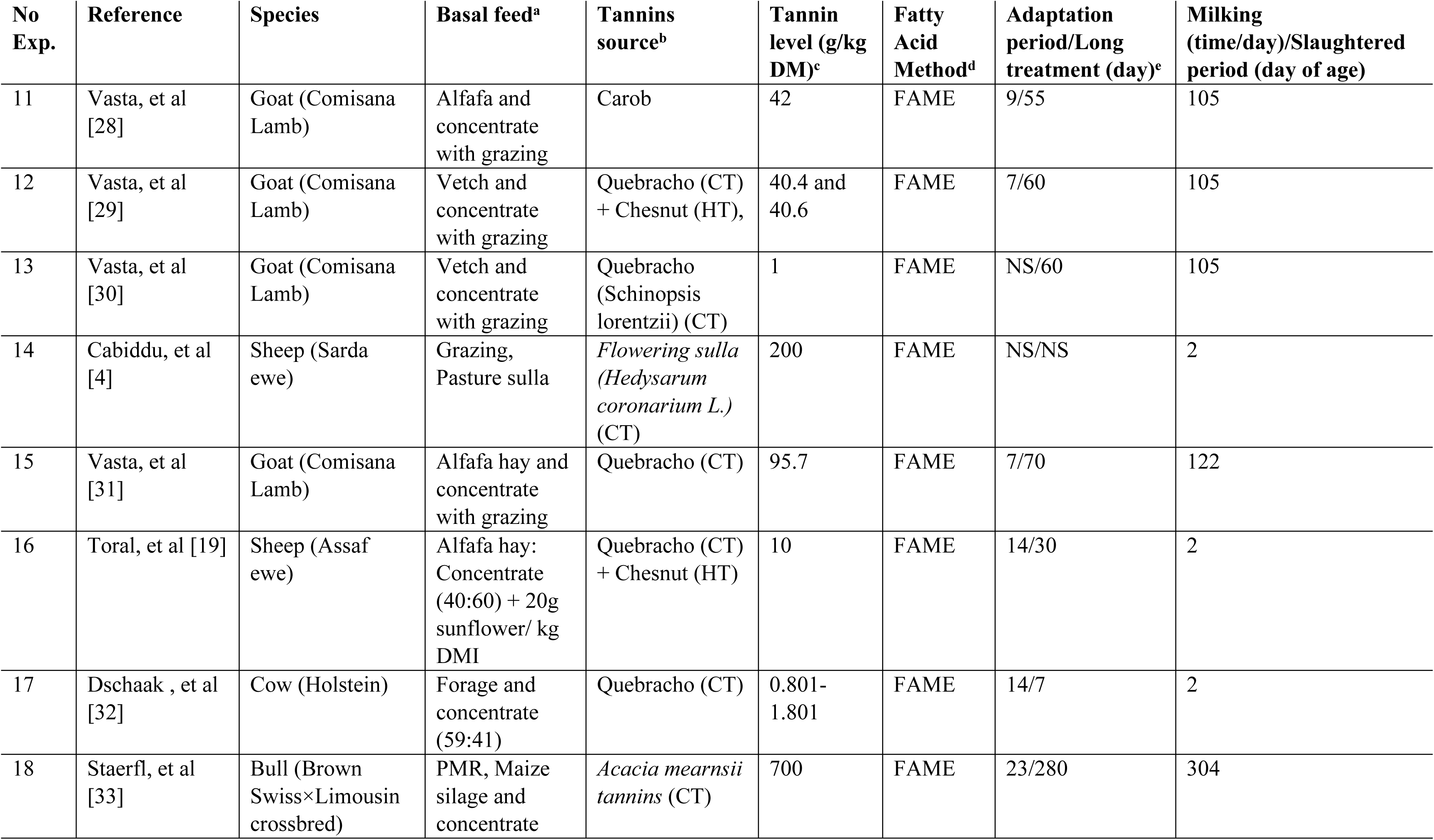
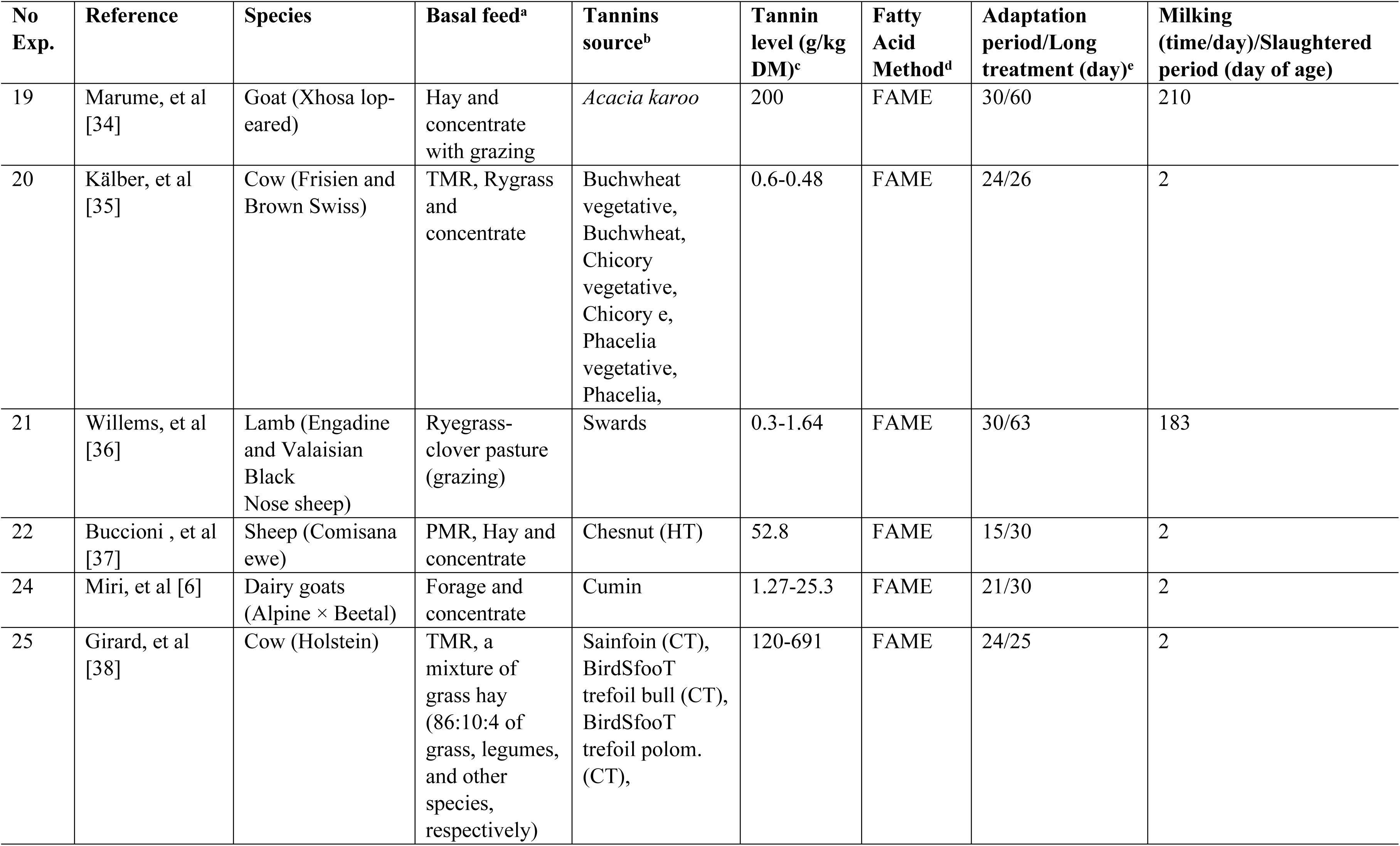
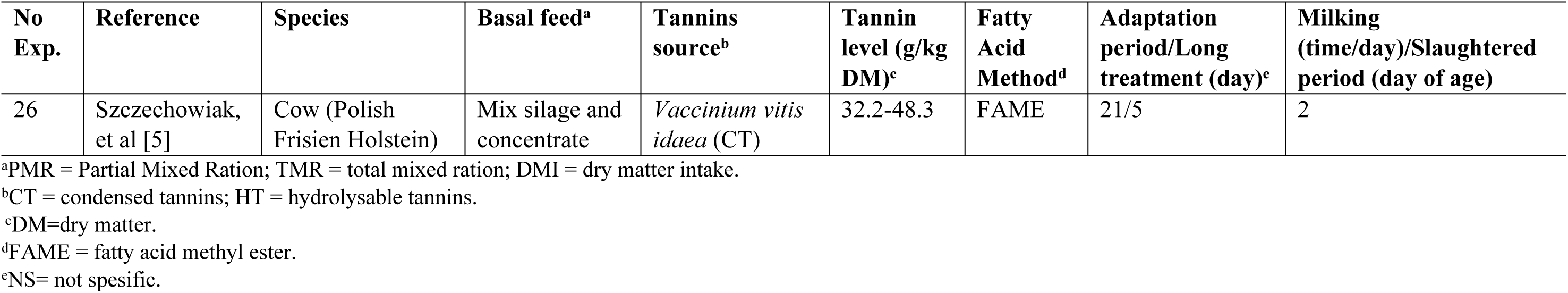
Data tabulation of *in vivo* experiments.

| No Exp. | Reference | Species | Basal feed<sup>a</sup> | Tannins source<sup>b</sup> | Tannin level (g/kg DM)<sup>c</sup> | Fatty Acid Method<sup>d</sup> | Adaptation period/Long treatment (day)<sup>e</sup> | Milking (time/day)/Slaughtered period (day of age) |
| --- | --- | --- | --- | --- | --- | --- | --- | --- |
| 11 | Vasta, et al [28] | Goat (Comisana Lamb) | Alfafa and concentrate with grazing | Carob | 42 | FAME | 9/55 | 105 |
| 12 | Vasta, et al [29] | Goat (Comisana Lamb) | Vetch and concentrate with grazing | Quebracho (CT) + Chesnut (HT), | 40.4 and 40.6 | FAME | 7/60 | 105 |
| 13 | Vasta, et al [30] | Goat (Comisana Lamb) | Vetch and concentrate with grazing | Quebracho (Schinopsis lorentzii) (CT) | 1 | FAME | NS/60 | 105 |
| 14 | Cabiddu, et al [4] | Sheep (Sarda ewe) | Grazing, Pasture sulla | *Flowering sulla* (*Hedysarum coronarium L.*) (CT) | 200 | FAME | NS/NS | 2 |
| 15 | Vasta, et al [31] | Goat (Comisana Lamb) | Alfafa hay and concentrate with grazing | Quebracho (CT) | 95.7 | FAME | 7/70 | 122 |
| 16 | Toral, et al [19] | Sheep (Assaf ewe) | Alfafa hay: Concentrate (40:60) + 20g sunflower/ kg DMI | Quebracho (CT) + Chesnut (HT) | 10 | FAME | 14/30 | 2 |
| 17 | Dschaak , et al [32] | Cow (Holstein) | Forage and concentrate (59:41) | Quebracho (CT) | 0.801-1.801 | FAME | 14/7 | 2 |
| 18 | Staerfl, et al [33] | Bull (Brown Swiss×Limousin crossbred) | PMR, Maize silage and concentrate | *Acacia mearnsii* tannins (CT) | 700 | FAME | 23/280 | 304 |
| 19 | Marume, et al [34] | Goat (Xhosa lop-eared) | Hay and concentrate with grazing | *Acacia karoo* | 200 | FAME | 30/60 | 210 |
| 20 | Kälber, et al [35] | Cow (Frisien and Brown Swiss) | TMR, Rygrass and concentrate | Buchwheat vegetative, Buchwheat, Chicory vegetative, Chicory e, Phacelia vegetative, Phacelia, | 0.6-0.48 | FAME | 24/26 | 2 |
| 21 | Willems, et al [36] | Lamb (Engadine and Valaisian Black Nose sheep) | Ryegrass-clover pasture (grazing) | Swards | 0.3-1.64 | FAME | 30/63 | 183 |
| 22 | Buccioni , et al [37] | Sheep (Comisana ewe) | PMR, Hay and concentrate | Chesnut (HT) | 52.8 | FAME | 15/30 | 2 |
| 24 | Miri, et al [6] | Dairy goats (Alpine × Beetal) | Forage and concentrate | Cumin | 1.27-25.3 | FAME | 21/30 | 2 |
| 25 | Girard, et al [38] | Cow (Holstein) | TMR, a mixture of grass hay (86:10:4 of grass, legumes, and other species, respectively) | Sainfoin (CT), BirdSfooT trefoil bull (CT), BirdSfooT trefoil polom. (CT), | 120-691 | FAME | 24/25 | 2 |

| No<br>Exp. | Reference | Species | Basal feed<sup>a</sup> | Tannins<br>source<sup>b</sup> | Tannin<br>level (g/kg<br>DM)<sup>c</sup> | Fatty<br>Acid<br>Method<sup>d</sup> | Adaptation<br>period/Long<br>treatment (day)<sup>e</sup> | Milking<br>(time/day)/Slaughtered<br>period (day of age) |
| --- | --- | --- | --- | --- | --- | --- | --- | --- |
| 26 | Szczechowiak,<br>et al [5] | Cow (Polish<br>Frisien Holstein) | Mix silage and<br>concentrate | *Vaccinium vitis*<br>*idaea* (CT) | 32.2-48.3 | FAME | 21/5 | 2 |

<sup>a</sup>PMR = Partial Mixed Ration; TMR = total mixed ration; DMI = dry matter intake.
<sup>b</sup>CT = condensed tannins; HT = hydrolysable tannins.
<sup>c</sup>DM=dry matter.
<sup>d</sup>FAME = fatty acid methyl ester.
<sup>e</sup>NS= not spesific.

### Statistical analysis

The analysis of the data assembled in the database was conducted by a statistical meta-analysis approach [13, 16, 17]. Using the MIXED procedure of SAS 9.4 version [18], the following model was applied:

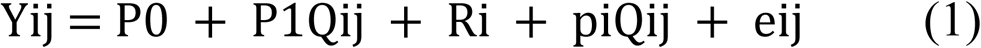

where Y_ij_ = dependent variable, P_0_ = overall intercept across all experiments (fixed effect), P_1_= linear regression coefficient of Y on X (fixed effect), Q_ij_ = value of the continuous predictor variable (supplementary tannin level), R_i_ = random effect of experiment _i_, pi = random effect of experiment i on the regression coefficient of Y on X in experiment i and e_ij_ = the deniable residual error. To input the CLASS statement, the variable ‘REFERENCENO’ was subjected without any quantitative information. Additionally, data were calculated by the number of animal replications in each experiment [18] and scaled to 1 to avoid misconception regarding unequal variance among experiments. In a fixed-effects model, a small study was considerably ignored, though, considerable weight was adjusted to a large study (based on number of measurements).

Outliers were identified by examining mixed procedure with maximum-likelihood (ML). For illustrating, it used METHOD=ML; COVTEST; PARMS statement followed by the EQCONS=2 option. An unstructured variance–covariance matrix (type = un) was confirmed as the random part of the model. Also, the comparison between CLA number from milk and meat source could not compare directly. It would be possible comparing total data including covering from *in vitro* and *in vivo* observations. Incompleteness of selected data on involving variables, meta analyses were technically performed based on the data available for individual variables.

## Results

### Search results and bias assessment

As depicted in Fig 1, identified articles had 11038 potentially relevant studies. Articles were checked compressing at 26 studies, see table 1 and 2. Twenty-six studies had performed the critically information, while 2 experiments conducted both in vitro and in vivo on the same time (Szczechowiak, et al [5] and Toral, et al [19]). According to Cochrane Reviewer’s Handbook 4.2 to assess risk of bias (S1 Fig). The high-risk studies were disclosed. Five of them had a low risk of bias.

### The effectiveness level of tannins

The meta regression strictly between dietary tannins and CLA levels from the *in vitro* batch culture experiment and the *in vivo* experiment is presented in Table 3 and Table 4, respectively. The optimum level of tannins for modulating CLA level coming along a nurture rumen fermentation was predicted around 0.1-50 g/kg DM. Regardless of tannin type, the tough natural chemists from tannins provoked the CLA going down gradually (p<0.001) of both studies in *in vivo* CLA milk shift (Fig 2) with an R^2^ of 0.929 and *in vivo* CLA meat precipitation (Fig 3) with an R^2^ of 0.549. However, supplementing a surge of tannin level increased the CLA level in *in vitro* study, yet, the efficiency of tannin acted dubious. Truly, a rising of CLA trend (p<0.001) was followed by a linear relationship rather than a quadratic response (Fig 4) with an R^2^ of 0.193.

**Table 3.**
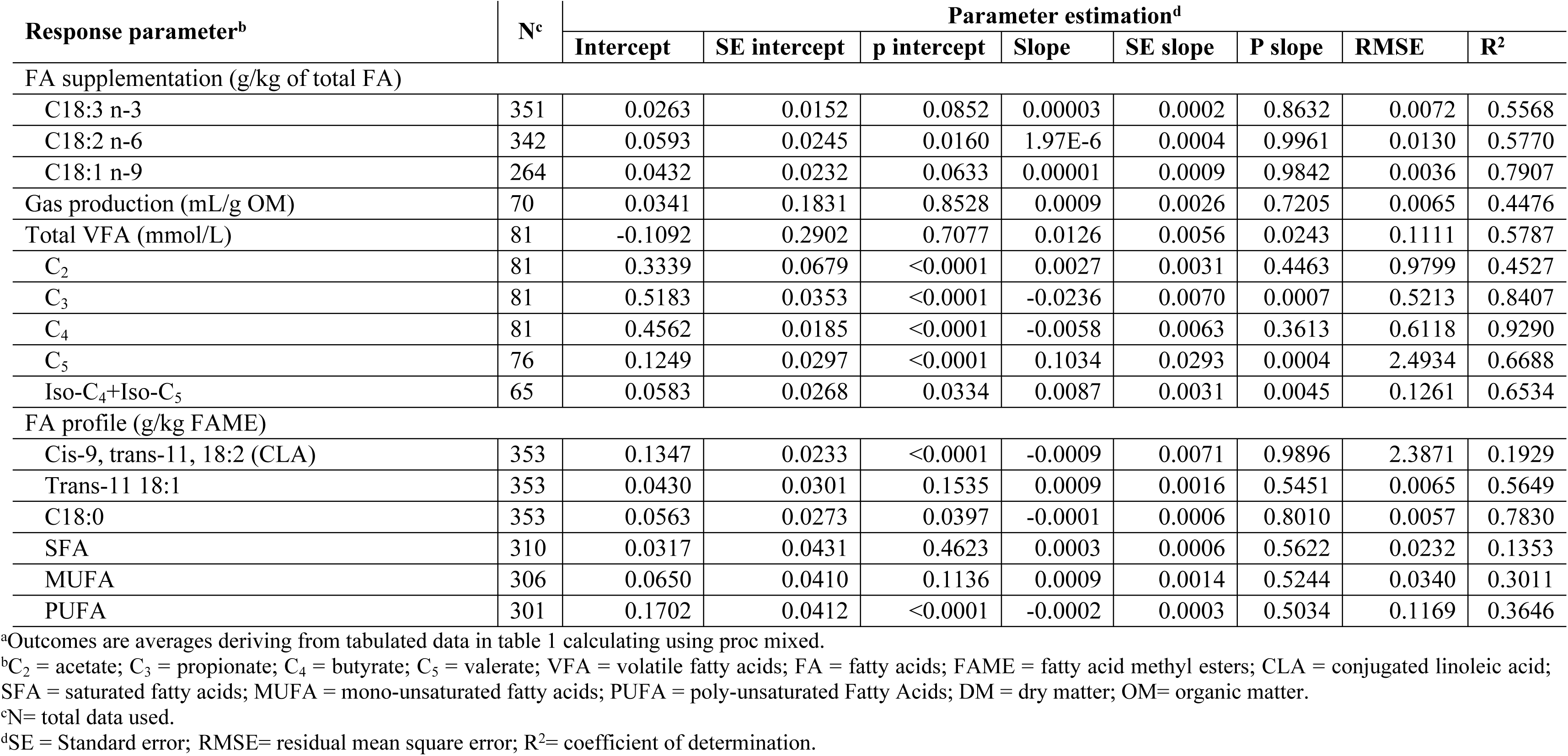
The predicting equation of *in vitro* batch culture experiments^a^.

**Table 4.** The predicting equation of *in vivo* batch culture experiments^a^.

| Response parameter<sup>b</sup> | N<sup>c</sup> | Parameter estimation<sup>d</sup> |  |  |  |  |  |  |  |
| --- | --- | --- | --- | --- | --- | --- | --- | --- | --- |
|  |  | Intercept | SE intercept | p intercept | Intercept | SE slope | P slope | Intercept | R<sup>2</sup> |
| FA supplementation (g/kg of total FA) |  |  |  |  |  |  |  |  |  |
| C18:3 n-3 | 276 | 0.3507 | 0.0328 | <0.0001 | -0.0040 | 0.0008 | <0.0001 | 0.8983 | 0.4374 |
| C18:2 n-6 | 278 | 0.2603 | 0.0206 | <0.0001 | -0.0071 | 0.0013 | <0.0001 | 0.5895 | 0.4519 |
| C18:1 n-9 | 247 | 0.2874 | 0.0438 | <0.0001 | -0.0093 | 0.0024 | <0.0001 | 0.7079 | 0.2651 |
| Total VFA (mmol/L) | 73 | 0.3462 | 0.4800 | 0.4731 | -0.0022 | 0.0050 | 0.6663 | 0.0192 | 0.4513 |
| C<sub>2</sub> | 73 | 0.0753 | 0.0428 | 0.0826 | 0.0007 | 0.0008 | 0.4166 | 0.0408 | 0.7222 |
| C<sub>3</sub> | 73 | 0.4125 | 0.0538 | <0.0001 | -0.0174 | 0.0051 | 0.0007 | 0.1785 | 0.7889 |
| C<sub>4</sub> | 73 | 0.4404 | 0.0621 | <0.0001 | -0.0338 | 0.0084 | <0.0001 | 1.4362 | 0.5928 |
| C<sub>5</sub> | 47 | 0.4218 | 0.0211 | <0.0001 | 0.0714 | 0.0316 | 0.0238 | 0.7399 | 0.9571 |
| Iso-C<sub>4</sub> + Iso-C<sub>5</sub> | 40 | 0.6673 | 0.0431 | <0.0001 | -0.1248 | 0.0212 | <0.0001 | 1.2507 | 0.8956 |
| Desaturation index |  |  |  |  |  |  |  |  |  |
| C18:2 cis-9 trans-11:C18:1 trans-11 | 60 | -0.2272 | 0.0191 | <0.0001 | 24.6657 | 1.5204 | <0.0001 | 4.7579 | 0.5538 |
| FA profile in milk (g/kg FAME) |  |  |  |  |  |  |  |  |  |
| Cis-9, trans-11 18:2 (CLA) | 281 | 0.1704 | 0.0129 | <0.0001 | 0.0967 | 0.0070 | <0.0001 | 1.6716 | 0.9289 |
| Trans-11 18:1 | 283 | 0.0676 | 0.0155 | <0.0001 | 0.0115 | 0.0027 | <0.0001 | 0.2682 | 0.3880 |
| C18:0 | 283 | 0.2181 | 0.0503 | <0.0001 | 0.0015 | 0.0016 | 0.3603 | 0.2653 | 0.3407 |
| SFA | 306 | 0.4411 | 0.0755 | <0.0001 | -0.0046 | 0.0013 | 0.0004 | 0.4115 | 0.3465 |
| MUFA | 295 | -0.0257 | 0.0577 | 0.6560 | 0.0083 | 0.0012 | <0.0001 | 0.4716 | 0.4552 |
| PUFA | 284 | 0.2370 | 0.0326 | <0.0001 | 0.0014 | 0.0008 | 0.0740 | 0.7109 | 0.4594 |
| FA profile in longissimus dorsi muscle (g/kg FAME) |  |  |  |  |  |  |  |  |  |
| Cis-9, trans-11, 18:2 (CLA) | 172 | 0.0193 | 0.0651 | 0.7682 | 0.0009 | 0.0330 | 0.9771 | 0.0026 | 0.5491 |
| Trans-11, 18:1 | 172 | 0.0102 | 0.0442 | 0.8183 | -0.0003 | 0.0068 | 0.9706 | 0.0005 | 0.4547 |
| C18:0 | 172 | -0.0035 | 0.0994 | 0.9720 | 0.0023 | 0.0053 | 0.6684 | 0.0026 | 0.3705 |
| SFA | 172 | 0.1035 | 0.1265 | 0.4146 | -0.0011 | 0.0031 | 0.7135 | 0.0088 | 0.3773 |
| MUFA | 172 | 0.0458 | 0.1912 | 0.8109 | 0.0004 | 0.0046 | 0.9270 | 0.0006 | 0.8220 |
| PUFA | 172 | 0.0517 | 0.1166 | 0.6583 | 0.0005 | 0.0032 | 0.8701 | 0.0006 | 0.8347 |

<sup>a</sup>Outcomes are averages deriving from tabulated data in table 1 calculating using proc mixed.
<sup>b</sup>C<sub>2</sub> = acetate; C<sub>3</sub> = propionate; C<sub>4</sub> = butyrate; C<sub>5</sub> = valerate; VFA = volatile fatty acids; FA = fatty acids; FAME = fatty acid methyl esters; CLA = conjugated linoleic acid; SFA = saturated fatty acids; MUFA = mono-unsaturated fatty acids; PUFA = poly-unsaturated Fatty Acids; DM = dry matter; OM= organic matter. <sup>c</sup>N= total data used. <sup>d</sup>SE = Standard error; RMSE= residual mean square error; R<sup>2</sup>= coefficient of determination.

**Fig 2.**
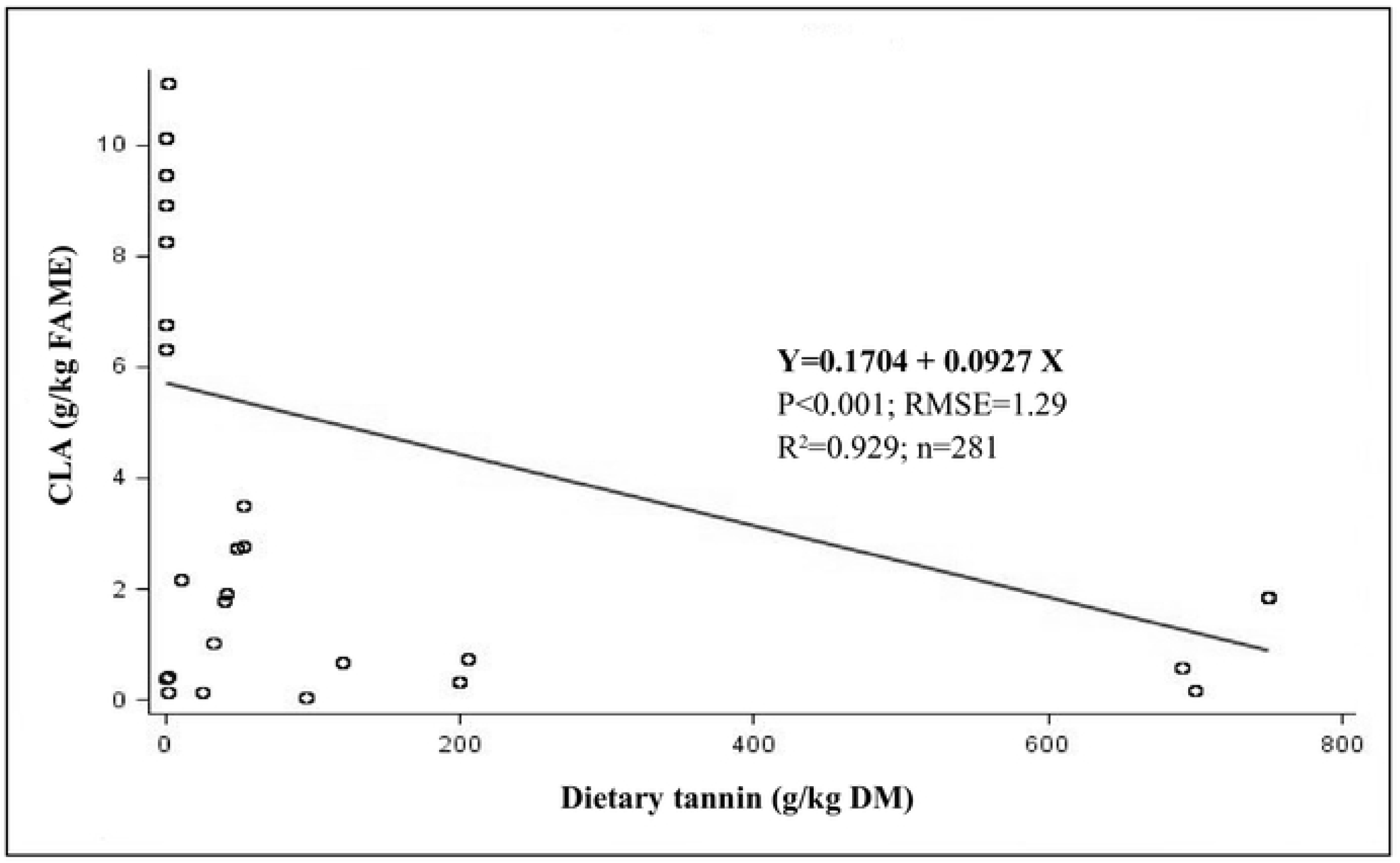
The linear relationship between dietary tannins (g/kg DM) and CLA milk shift (g/kg FAME) using *in vivo*.

**Fig 3.**
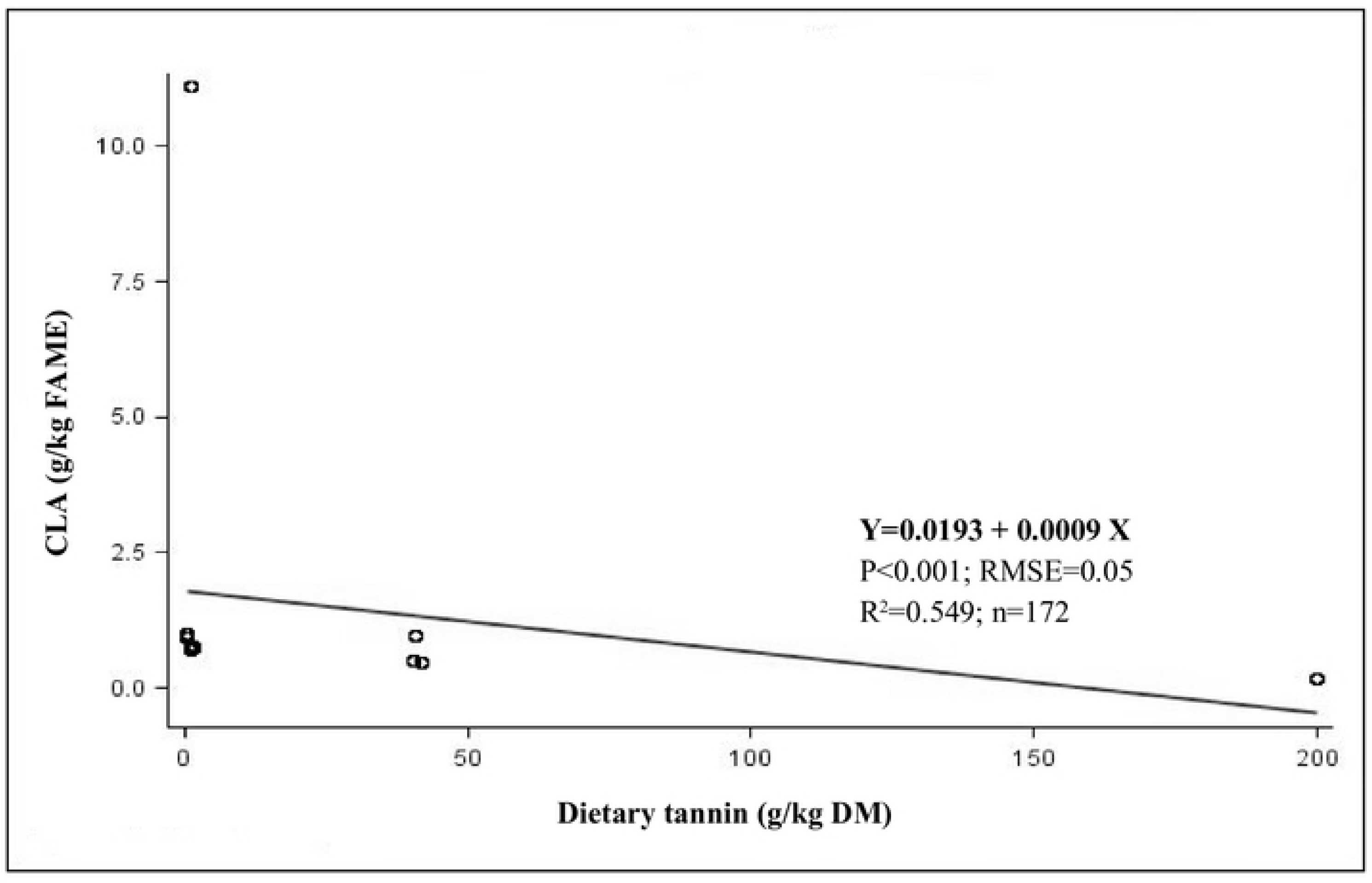
The linear relationship between dietary tannins (g/kg DM) and CLA meat shift (g/kg FAME) using *in vivo*.

**Fig 4.**
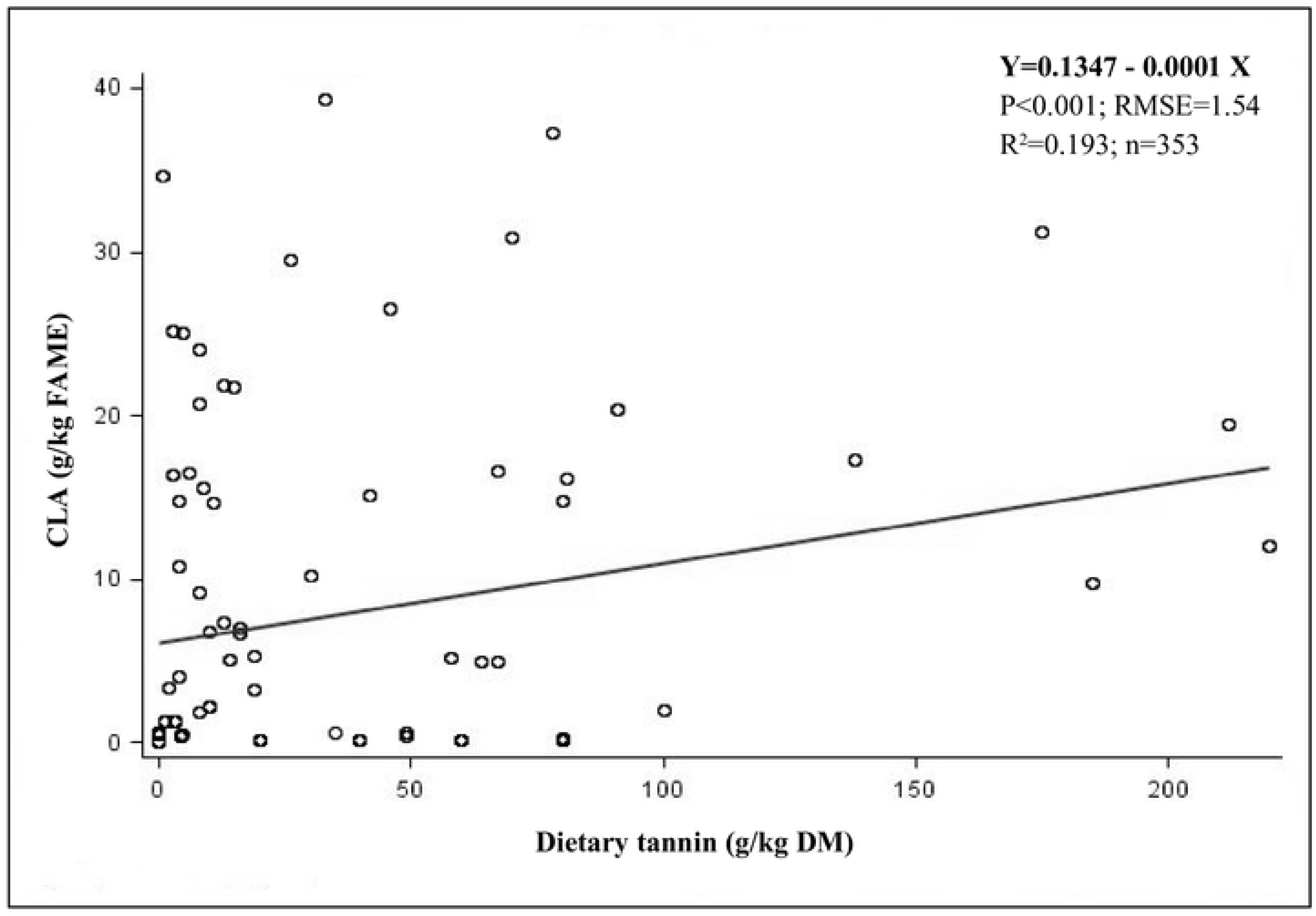
The linear relationship between dietary tannins (g/kg DM) and CLA (g/kg FAME) using *in vitro*.

### The regression of method application

The regression relationships between the *in vitro*-*in vivo* of CLA milk form is depicted in Fig 5 and *in vitro*-*in vivo* of CLA meat deposition shown in Fig 6. These relationships were expressed by a linear relationship rather than a quadratic response. Clearly, there were no relationship among them (R^2^<0.1).

**Fig 5.**
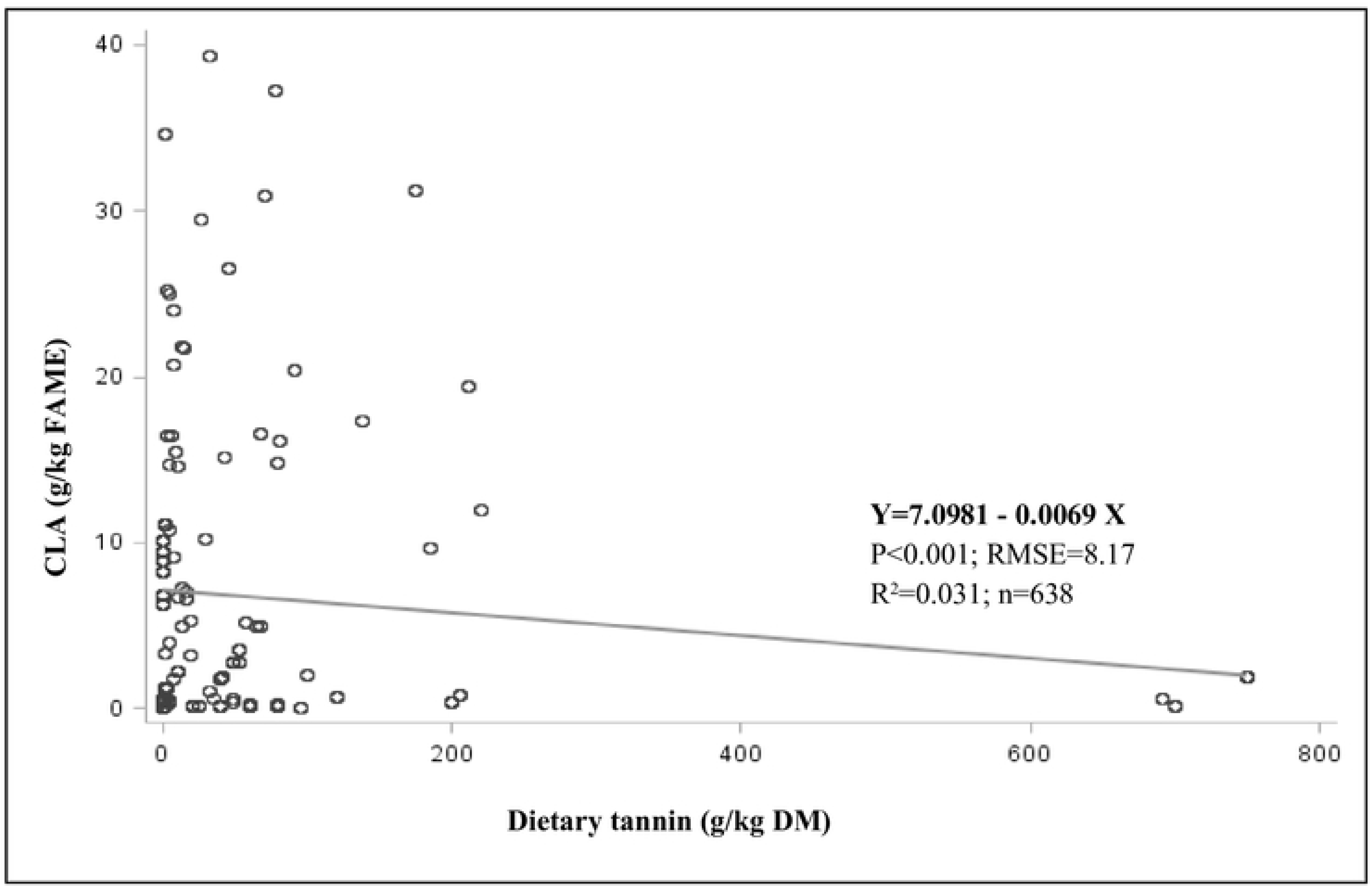
The relationship between *in vitro* and *in vivo* CLA milk form.

**Fig 6.**
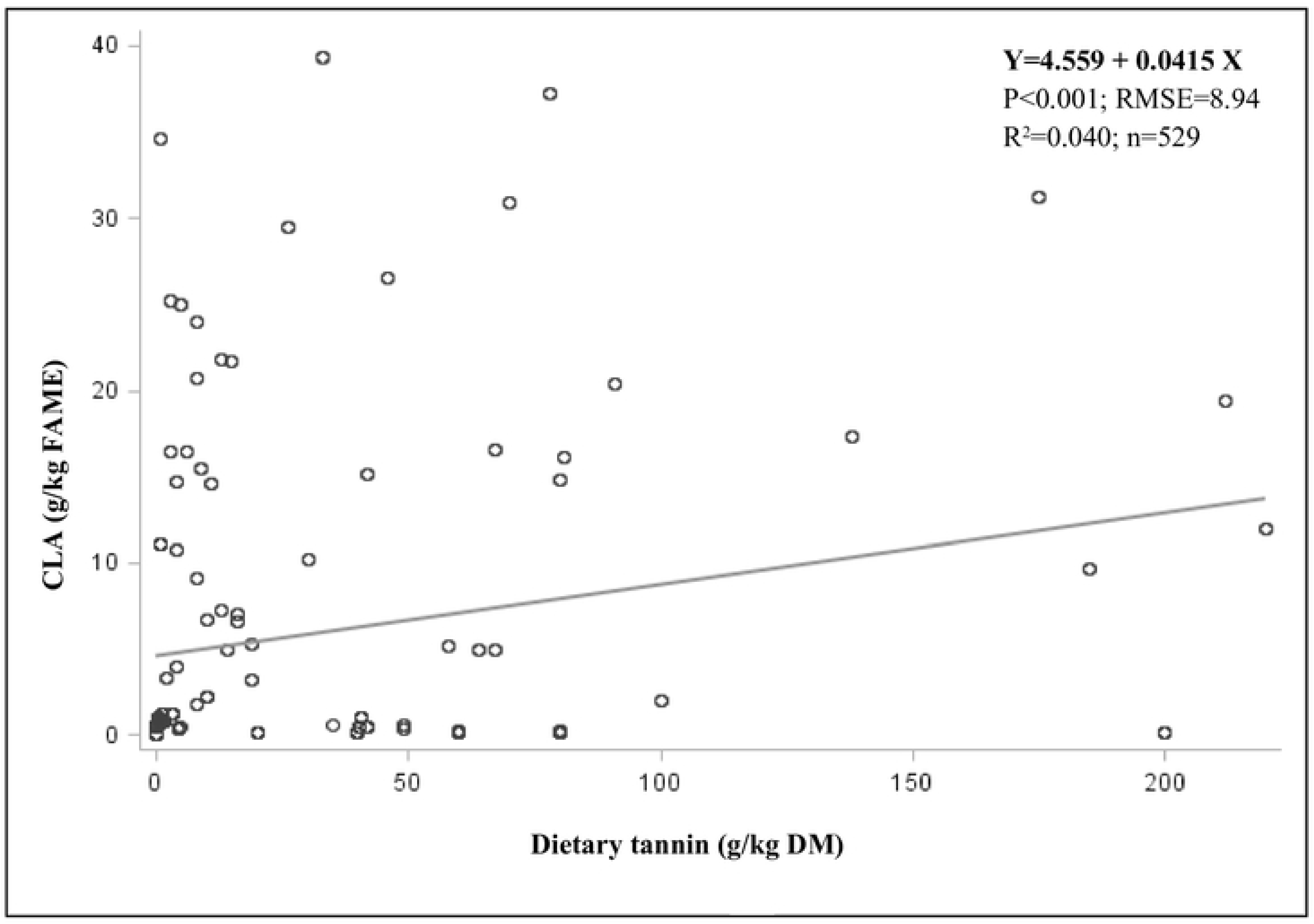
The relationship between *in vitro* and *in vivo* CLA meat form.

## Discussion

In this meta-analysis provided underlying prediction and perspective in other tannin ways such a manipulating biohydrogenation to compulsive CLA production massively. As shown in Table 3, dietary tannin affected as statistically regression of the CLA components in milk form (R^2^>0.9), however, not in meat precipitation (R^2^<0.9) without collecting bias data. These results were indirectly as same as earlier meta-analysis predicting dietary tannin level on rumen diet^12^. Enhancing effort to modulate biohydrogenation is rough and tricky, wherein choose a suitable method to approve. More elaborating reasons, addressed by Lourenço, et al [1], a role for manipulating biohydrogenation was difficult because of inviting systematically isomerization through decreasing hydrogen supply and assessing stearic (C18:0) bacteria. Unfortunately, this meta-analysis was at dull in rumen bacterial biohydrogenation in order to limitation of public-access records.

One way to obtain information of this study was merely understanding to input a sufficiency feedstuff, particularly fiber and fatty acid source, to start fermented nutrient in creating gas production including hydrogen accumulation that could be as references continuously on biohydrogenation. Castro, et al [39], shown diet containing more fat source increased desaturation index, though there was lower regression number of this regard (R^2^<0.9), see table 4. In addition, whether in *in vitro* and/or *in vivo* demonstrated low regression (R^2^<0.9) of fatty acid role on predicting tannin properties, see table 3 and 4. Another, trans 11 C18:1 (vaccenic acid) were not affected (p<0.001, R^2^<0.9) by dietary tannin in all methods. Hence, there was a relationship between FA supplementation with desaturase index on ruminal biohydrogenation.

Furthermore, Jayanegara, et al [13], the regression of tannin effect on rumen metabolism and its methane loss provoked a lesser biohydrogenation indirectly, through diminishing a hydrogen (H_2_) supply and contribution of volatile fatty acid (VFA’s). This was corresponding to a higher propionate catching H_2_ down leading to greater biohydrogenation failure. In this meta-analysis, the propionate was interrupted (p<0.001) by tannin supplementation with a tantamount R^2^value. In same way, Dschaak, et al [32], reported supplementary condensed tannin in different forage levels interpreted an increase of propionate, yet, no affection for distributed CLA. However, Buccioni, et al [37], declining propionate at 34.3% against control diet tending to increase of CLA formation around 24.2% in milk production, when dairy ewes fed quebracho as tannin-containing feedstuff. It might be tannin form inducing the different sub-active compound inside leading to the different affections and this mechanism would be only running on fat metabolism persuading a recycling lively organ such a liver. Although, *in vitro* study reported in different way in this study, see table 3.

Comparison of selectively differential methods by present of CLA fractions are possibly corresponded each other with similar units of measurement. To be recognized, the *in vitro* bath culture method ran dissimilar towards to the *in vivo* method in this study. Nevertheless, the media flow out substantially carried on their metabolism and there was lively absorption of the rumen properties directly on the process. Remember, in this meta-analysis concern, one unit was presenting on the graph by CLA (g/kg FAME) to clarify the relationship between CLA *in vitro* relating to CLA milk *in vivo* (Fig 5) and CLA *in vitro* relating to CLA meat *in vivo* (Fig 6). Astoundingly, both of their relationships had expressed as to be poor regression (R^2^<0.5) [40]. Thus, it could be clarified that being challenging to predict from *in vitro* observation to *in vivo* situations accurately on CLA property determination of the ruminants^5,40,^and/or field objective close to FA measurement^41^. It was known in advance that the *in vitro* observation presenting with a current limitation, especially to extrapolate how systematically synthesizing biohydrogenation of fatty acid was.

Utterly, dietary tannins supplementation to ruminants sentenced multiparous benefits, especially at CLA production. The most significant findings in this study were that the ruminants achieving tannin-containing diet altered rumen fermentation leading to direct and indirect effects on biohydrogenation. Yet, the suitable method was considered as perquisite trial whether *in vitro* and *in vivo* studies. Therefore, dietary tannin may change other specifically parameters, for instance behavior of gene expressions for further investigations needed.

## Conclusion

Coming with sizeable data from the valid publications, this meta-analysis provided a prediction of suitable plant-containing tannin level in ruminal diet and their application facing a fit method design for developing CLA formation on biohydrogenation. The optimum level of tannins was predicted around 0.1-5.0 g/kg DM. Basically, adjusting level of tannins declined the CLA number. Secondly, the *in vivo* method was more suitable for directly observation that concerned in FA transformation. Unless, using the *in vitro* observation was easier, cheaper, and more edible presenting with a current limitation, particularly to understand the full outcome of systematically synthesizing FA on biohydrogenation.

## Supporting information

**S1 Table. Full electronic search strategy for ISI Web of Knowledge.**

**S1 Fig. Bias assessment using Cochrane Reviewer’s Handbook 4.2.**

**S1 Checklist. PRISMA checklist.**

## Acknowledgment

Authors would like to say thanks to Laurence V. Madden, Amonrat Molee, and Jan Thomas Schonewille for preparing database design.

## Author contribution

### Conceptualization, methodology, investigation, resources, data curation and writing— original draft preparatio

Rayudika Aprilia Patindra Purba.

### Funding acquisition, project administration and writing—review and editing

Rayudika Aprilia Patindra Purba, Pramote Paengkoum, Siwaporn Paengkoum.

